# Shading does not lower thermal tolerance in the coral *Montipora capitata*: implications for conservation intervention

**DOI:** 10.1101/2024.11.13.623387

**Authors:** Hugo Ducret, Christopher R. Suchocki, Claire E. Bardin, Claire J. Lewis, Tristan Permentier, Madeleine Hardt, Robert J. Toonen, Marc Kochzius, Jean-François Flot

## Abstract

Marine heatwaves trigger severe coral bleaching events that result in dramatic losses of coral reefs worldwide. An increasingly common method used to mitigate coral bleaching is to shade portions of reefs. However, as shaded corals become less exposed to environmental stress, shading has also been hypothesized to lower their thermal tolerance, which would be detrimental for their survival. Here, we investigated how long-term shading modifies the response of *Montipora capitata* to both light and temperature stressors. After two years of growth, we used the Coral Bleaching Automated Stress System (CBASS) to compare responses of 73% shaded and unshaded corals during 1) temperature stress under ambient light, 2) temperature stress under low light, and 3) ambient temperature with light stress. Results show that shaded corals are less resistant than control ones to light stress and to heat stress under high light, but not to heat stress under low light, where no difference was detected. We further demonstrate that low light intensity during heat stress reduces the decline of photosynthetic efficiency when compared to heat stress at ambient light, regardless of the light history of the coral colonies. Our data support light-mediated stress, both independent of and synergistic with heat-induced stress, and suggest that light and heat stresses trigger different molecular and cellular pathways. We hereby confirm the benefits of coral shading under both short– and long-term temperature stress, and underline the importance of light acclimation in the conservation and restoration of coral reefs.

## Introduction

Coral reefs are threatened by ocean warming worldwide and are experiencing increasingly severe pantropical thermal anomalies. These phenomena trigger coral bleaching, a breakdown of the coral-algal symbiosis that results in large-scale coral mortality and degradation of coral reef communities (Hughes et al. 2017). Bleaching events are predicted to increase in frequency and intensity as ocean temperatures rise in the coming decades. Model predictions indicate that more than 75% of the world’s coral reefs may experience annual severe bleaching before the end of the century (van Hooidonk et al. 2016; Evensen et al. 2022), which would have dramatic ecological, economical, and human consequences (Moberg & Foke 1999).

In this context, understanding the exact causes and mechanisms of coral bleaching is of paramount importance to devise conservation interventions (Bowgen et al. 2022). Bleaching can be induced by a wide variety of abiotic drivers and metabolic pathways due to different sensitivities of host and symbiont cells (Oakley & Davy 2018; Helgoe et al. 2024). Among the main stressors, excesses of light and high temperature are known to have independent but often synergistic effects on the induction of bleaching (Coles & Jokiel 1978; Helgoe et al. 2024). Irradiance levels beyond tolerance thresholds usually result in protein damage in the symbionts’ Photosystem II, which stops the production of NADPH and ATP, thus decreasing the electron transport rate between photosystems (Weis 2008; Roberty et al. 2014 & 2015; Helgoe et al. 2024). Such imbalance between donors and acceptors leads to the formation of Reactive Oxygen Species (ROSs) in the symbiont’s cells, which can diffuse to the host’s cells and cause oxidative damage there (Baird et al. 2009; Lesser et al. 2011). The overwhelming concentration of ROSs in the host’s cells is responsible for initiating a caspase cascade, eventually leading to apoptosis (Weis 2008; Baird et al. 2009; Paxton et al. 2013; Oakley et al. 2017, Oakley & Davy 2018; Helgoe et al. 2024), wherein symbionts are expelled and coral skeleton becomes visible. Alternatively, bleaching can also be triggered by an increase of temperature in complete absence of light (Tolleter et al. 2013). High water temperature drives protein misfolding in the host’s endoplasmic reticulum (ER, Orrenius et al. 2003; Oakley & Davy 2018), as well as high respiration leading to an inhibition of the electron transport rate in the host’s mitochondria (Nii & Muscatine 1997; Weis 2008; Baird et al. 2009) and damage to the symbiont’s chloroplasts (Warner et al. 1999; Takahashi et al. 2004). Eventually, these different processes also involve the massive production of ROS by-products, leading to multiple metabolic dysfunctions in both the host and the symbiont, and eventually to necrosis or apoptosis (Helgoe et al. 2024). This example of different triggers and cascades highlights the complexity of coral bleaching and the importance of designing context-specific solutions to mitigate it.

A growing number of approaches are being developed to tackle these environmental stressors and minimize coral loss (Albright & Cooley 2019; Anthony et al. 2020; Voolstra et al. 2021). One of these practices is derived from the observation that bleaching susceptibility decreases with depth (Baird et al. 2016; Loya et al. 2017; Bongaerts & Smith 2019; Winslow et al. 2024), as well as in other naturally occurring conditions that reduce light intensity, such as increased turbidity (van Woesik & al. 2012), crevices and overhangs (Hoogenboom et al. 2017), or increased cloud cover (Mumby et al. 2001, Gonzalez-Espinosa & Donner 2021). Reduced light intensity decreases light stress, which lowers the synergistic impact of light and temperature on essential metabolic pathways (Roth 2014). Therefore, there has been a growing interest in implementing and studying the impact of shading on coral reefs (Coelho et al. 2017, Tagliafico et al. 2022; Butcherine et al. 2023). This strategy would target small, genetically diverse portions of reefs, or populations that could provide a source of propagules to rapidly reseed a bleached reef and mitigate extinctions, although attempts to expand it to a larger scale are underway (Tollefson 2021; Tagliafico et al. 2022). However, as shaded corals become less exposed to environmental stress, decreased light intensity could actually lower stress tolerance, which would be detrimental to their survival. Corals can acquire resistance to heat-induced bleaching via prior exposure to high solar radiation (Brown et al. 2002; Schoepf et al. 2015) or to highly fluctuating, naturally extreme temperature environments (Schoepf et al. 2015). Alternatively, shaded corals acclimatize to a darker light regime which may reduce their exposure to stress over the long term, such that removal of shading may actually have deleterious consequences.

More generally, the extent to which light acclimation impacts the biology of coral holobionts remains poorly understood (Iluz & Dubinsky 2015). While thermal thresholds have been extensively studied across species and locations (Howells et al. 2012; Kumagai et al. 2018; Dilworth et al. 2021; Evensen et al. 2022), light thresholds, or light niches, remain under-studied, although there is interest in the related concept of depth niches (Roberts et al. 2019; Montgomery et al. 2021). Solar irradiance decreases exponentially within the water column, with site-specific variations (Kahng et al. 2019), and therefore depth niches can act as a useful proxy for light tolerance (Roberts et al. 2019; Montgomery et al. 2021). Most species are depth– and therefore light-specialists with relatively narrow depth preferences, but there are species that show extraordinary range in their depth distribution and therefore assumably in their physiology and photobiology (Roberts et al. 2019; Padilla-Gamiño et al. 2019; Montgomery et al. 2021). This applies to the Hawaiian rice coral *Montipora capitata*, which can be found from 1 to 100 meters deep (Veron 2000). Within a conservation framework, reliable estimation of light tolerance would provide critical information on optimal outplant site, as well as whether a period of light acclimation may be required prior to moving corals from a higher-light area. Pathways and endpoints of coral bleaching involving the combination of light and heat may also be expected to differ between shallow and deeper colonies of a same species (Helgoe et al. 2024; Winslow et al. 2024), or of the same fragmented individual. Lastly, disentangling heat stress, light stress, acclimation, and bleaching is a critical step in our understanding of coral responses to global climate change (Gomez-Campo et al. 2022).

In this study, we investigated whether long-term exposure to decreased light intensity impacts the stress response of *Montipora capitata* corals. At the Hawai‘i Institute of Marine Biology (HIMB), we conducted a two-year, in-situ shading experiment (Fig. 1) wherein we grew coral fragments in the waters surrounding Moku o Lo‘e. At the conclusion of this experiment, we compared the performance of shaded and unshaded *M. capitata* to temperature stress under ambient light, temperature stress under low light, and light stress at constant water temperature.

**Fig. 1:**
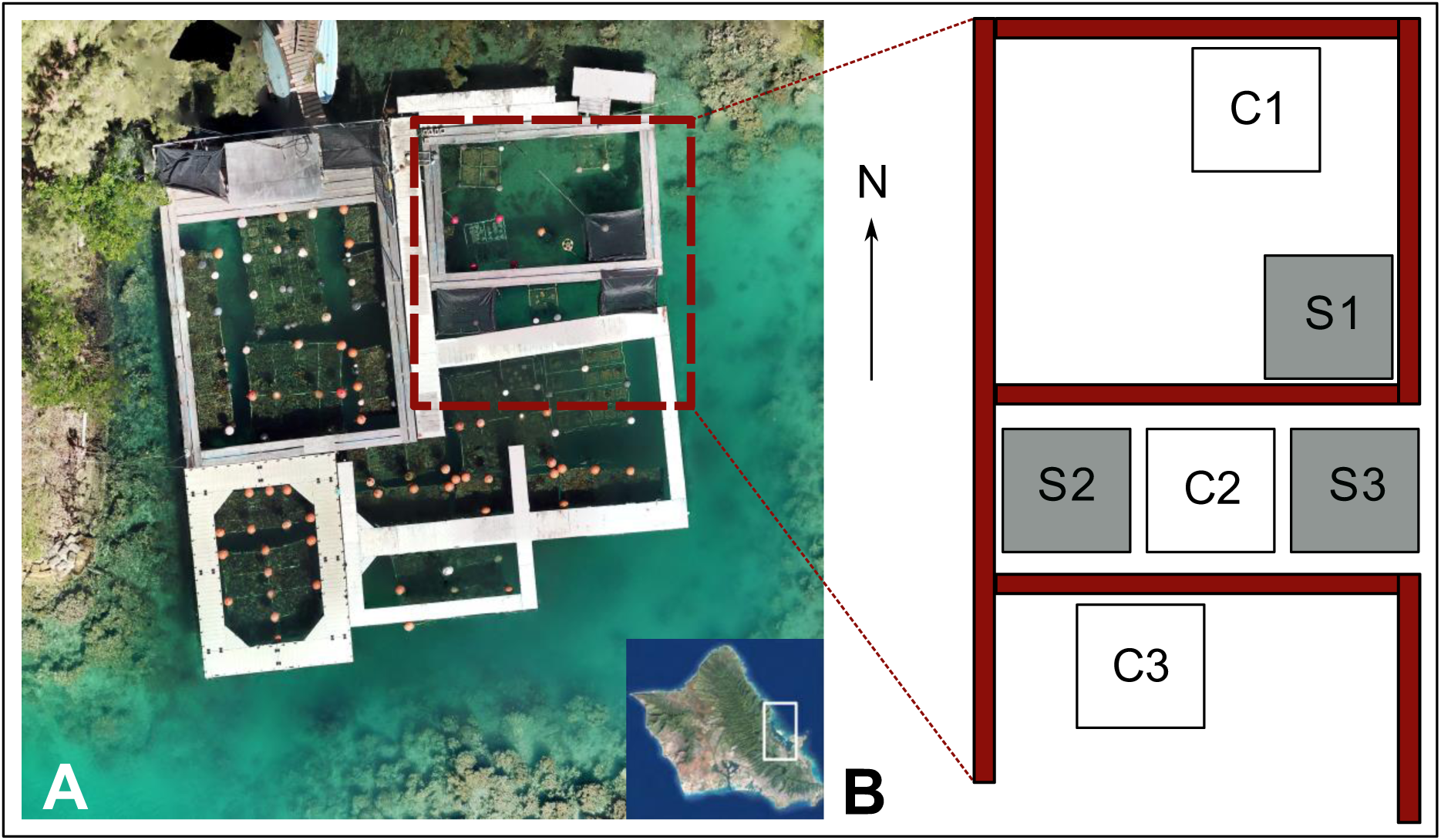
A) Aerial view of the Hawai’i Institute of Marine Biology coral nursery. The racks used in this study are located in the top-right corner (i.e. in the frame). The map of O‘ahu was retrieved from Google, with the white frame showing the location of Kāneʻohe Bay. B) Schematic representation of the experimental design, placed on the eastern side of the nursery. White squares represent control racks (C1,C2,C3), while gray boxes depict racks upon which a 73% shade cloth was deployed (S1,S2,S3).

## Methods

### Experimental design

The HIMB coral nursery consists of floating walkways surrounding and supporting suspended midwater platforms – or “racks” – for coral cultivation (Fig. 1, Knapp et al. 2022). In our experiment, six experimental racks were built at 2m depth, which corresponds to 1) the natural occurrence of *M. capitata* in the neighboring reef and 2) the depth at which corals typically grow in the nursery. Three of the racks were unshaded and exposed to ambient light while the other three were shaded (Fig. 1) using 6’ x 8’ black mesh screens with 73% shading efficiency (EZ corners, O‘ahu, Hawai‘i). We placed the fabric approximately 15 cm above the water surface. Additionally, shade cloth was added on the sides to shade corals from morning and evening irradiance. We randomized control and shade racks in the nursery to avoid any locality effects. HOBO Pendant data loggers (Onset, USA) were placed on each rack to record light intensity and water temperature in five-minutes intervals. Subsequently, nine large *M. capitata* coral colonies of about 40 cm in diameter chosen randomly from the nursery were fragmented into six pieces (n = 54). The corals in the nursery have naturally recruited to the racks over time or have grown out from previous leftover fragments and do not need any permit for collection or use. Each rack received a piece from each colony, which resulted in nine fragments per rack, or twenty-seven fragments in each treatment. Fragments were positioned randomly on each rack to avoid any locality or orientation effect. After fragmentation, corals were left to recover for two weeks at ambient light conditions before shade was applied. Top-down photographs of each colony were taken at t_0_ and every six months to monitor growth and mortality.

### Physiological measurements

In order to gain insight into how light intensity impacted coral photophysiology during the two years experiment at the nursery, we measured key physiological properties such as photosynthetic efficiency, symbiont density, and growth rate.

Prior to shading, corals’ steady-state photosynthetic efficiency (ΔF/F_m_’) was assessed using a diving-PAM (WALZ, Germany). This ratio, also referred to as effective quantum yield, is an indication of the amount of energy used by Photosystem II (PSII) under steady-state photosynthetic lighting conditions (Lichtenthaler et al. 2005). The higher the ratio, the higher the photosynthetic rate. Once shade clothes were applied, corals’ ΔF/F_m_’ was monitored daily for one week. All measurements were taken at noon on vertically oriented tips (which are the most exposed to sunlight) to limit variation.

As corals rely upon their photosynthetic symbiotic algae to meet their daily metabolic energy requirements, the density of symbionts is known to vary depending on the amount of light available in the environment (Wall et al. 2020). To test for differences in symbiont densities among treatments, we measured symbiont density after six months of growth following Wall et al. (2020). After coral tissues were removed from the skeleton using an airbrush with filtered seawater (0.2µm), the tissue slurry was then homogenized, and aliquots were taken for physiological measurements. Symbiont cell densities were determined using an automated cell counter (Countess 3, Thermo Fisher Scientific) and were standardized to the surface area of the coral fragments (Edmunds & Gates 2002).

Lastly, corals were weighed using the buoyant weight method (Jokiel et al. 1978). Each individual was weighed at t_0_ and again six months later in order to measure relative growth rates. As small fragments present a higher surface-area-to-volume ratio than large ones, their relative growth rate is usually higher than those of larger fragments (Soper et al. 2022). Therefore, growth rate was standardized to buoyant weight at t_0_ (Fig. 2D).

**Fig. 2:**
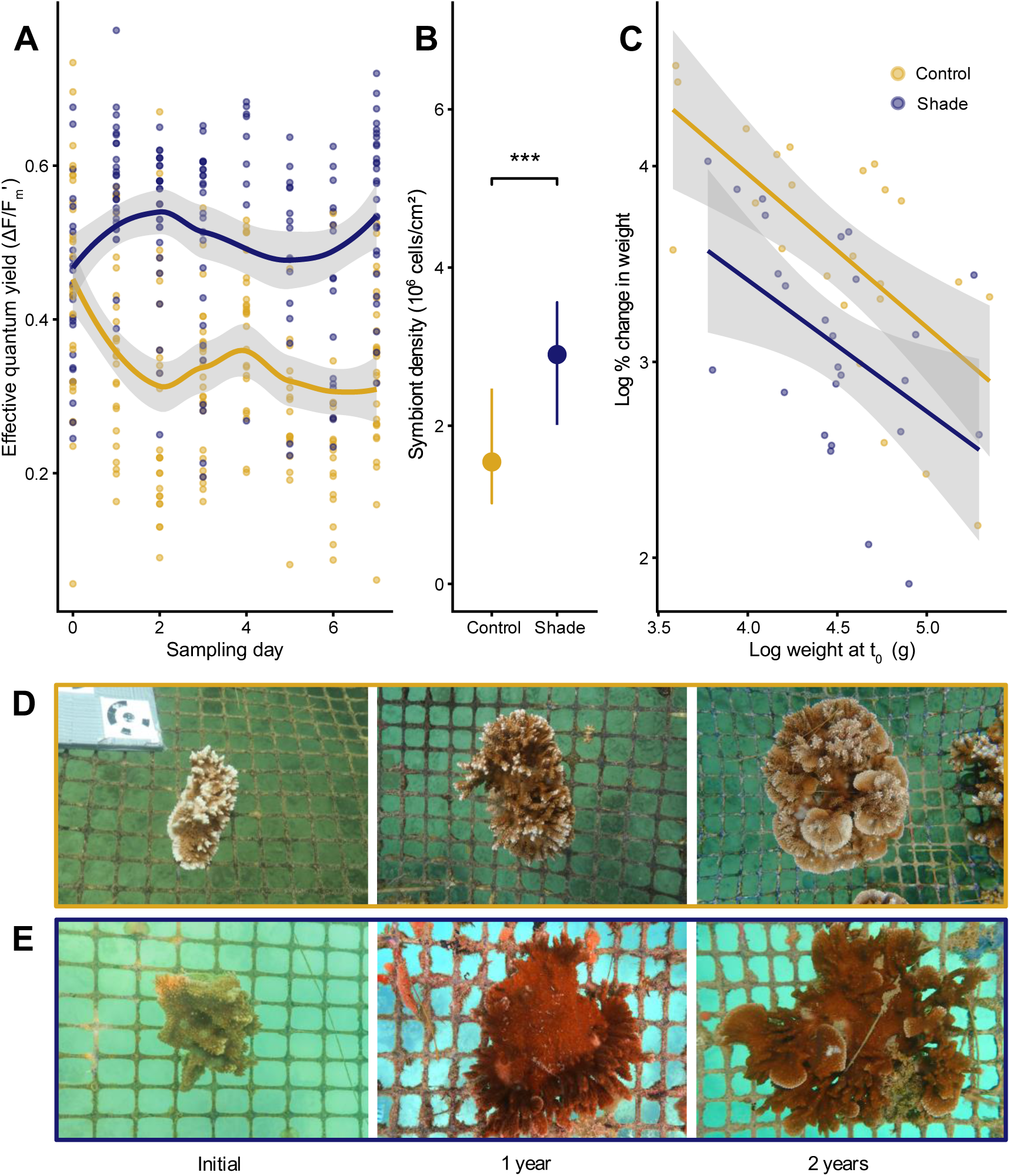
A) Photosynthetic efficiency time-series of corals taken during the first days of the shading experiment at the coral nursery. Curves show the main ΔF/F_m_’ value for each treatment and each sampling day, and gray areas depict standard deviation. B) Symbiont density pointranges of shaded and unshaded corals after a year and a half of experiment. C) Expression of the log-transformed growth rate against log-transformed initial weight at t_0_ for corals of each treatment over six months. The log-log scale was used in order to obtain a linear – i.e. more visual – relationship. Solid lines depict the main trend obtained from a linear regression between these two transformed variables, while gray areas represent standard deviations. Below are time-series of two coral fragments originating from the same colony, that grew at ambient (D) and 73% shaded (E) light, and from which nubbins were cut to perform the three CBASS assays after two years of growth

### Coral Bleaching Automated Stress System (CBASS) assays

After two years of growth at ambient or reduced light in the coral nursery, we used the Coral Bleaching Automated Stress System (CBASS) to compare stress responses of shaded and unshaded *M. capitata* fragments (Evensen et al. 2023). The CBASS set-up consists of a set of four 10-L flow-through tanks, in which conditions range from no heat or light stress in the first tank to very acute heat or light stress in the last tank. Each tank comprise two 200W titanium heaters (Bulk Reef Supply, USA), four 50W thermoelectric chillers (Nova Tec, USA), two 2000L.h^-1^ powerheads (SunSun, China), and a 200W full-spectrum LED light (Phlizon, China). Heaters and chillers are controlled through custom-built controllers based on the open-source platform Arduino, which allow for a reliable, fine-scale control of the seawater temperature. CBASS makes it possible to analyze the biological response of corals to site-specific temperature profiles at a user-defined light intensity level. In addition, each CBASS assay lasts 8 hours, which allows for repeated experiments within a small time frame, avoiding biases caused by environmental fluctuations and differences linked with seasonality. For each CBASS assay, we selected five *M. capitata* genotypes from each treatment and placed three replicates of each genotype per tank, for a total of n = 120 fragments per CBASS assay (Evensen et al. 2022). All coral nubbins were cut twenty-four hours prior to CBASS, and randomized within each tank. We used nubbins from the same ten coral pieces (five genotypes from each treatment) across all CBASS assays to avoid group effects between each assay.

Detailed temperature and light profiles for each tank are shown in Supplementary Fig. 2. Each eight-hour assay started at 1pm. The assay begins with a three-hour ramp to the targeted dose (i.e. increased temperature or light) followed by a three-hour hold at their intended treatment, and then finally they ramp back to their initial conditions over one hour. All tanks received flow-through seawater at a rate of ∼2L per hour.

The first CBASS assay consisted of a temperature stress under ambient, constant light intensity (600µmol.m^-^².sec^-1^), as specified in the CBASS standard protocol. The four temperature treatments were as follows (Fig. 3); 27°C (Kāneʻohe Bay Maximum Monthly Mean, MMM), 30°C, 33°C, and 36°C, as outlined in Evensen et al. (2023). For the second CBASS assay, we used the same temperature profiles, but lowered the light intensity to a constant 70µmol.m^-^².sec^-1^ to mimic the light intensity received by corals from the shade treatment (Supplementary Fig S1). In the last CBASS assay, we exposed the nubbins to differing light stress at a constant water temperature of 27°C (MMM) in all tanks. A profile similar to the temperature profiles was used for this light stress assay, with a three-hour ramp to targeted light intensity, a three-hour hold, and a one-hour ramp down. Tank 1 remained at a constant light regime of 70µmol.m^-^².sec^-1^ and a 510µmol.m^-^².sec^-1^ increase was applied to each subsequent tank. The resulting targeted irradiances were 70µmol.m^-^².sec^-1^, 580µmol.m^-^ ².sec^-1^, 1090µmol.m^-^².sec^-1^,and 1600µmol.m^-^².sec^-1^ for each of the four tanks respectively. The targeted light intensity in Tank 2 corresponds to the level of irradiance received by coral nubbins during the first temperature assay, while light profiles in Tank 3 and 4 were designed to be acute light stressors. After the three-hours hold at their target light intensities, lights were all switched off at 7pm. For this assay, light intensity was also measured by HOBO pendant data loggers (Onset, USA) with measures taken on a five-minutes interval.

**Fig. 3:**
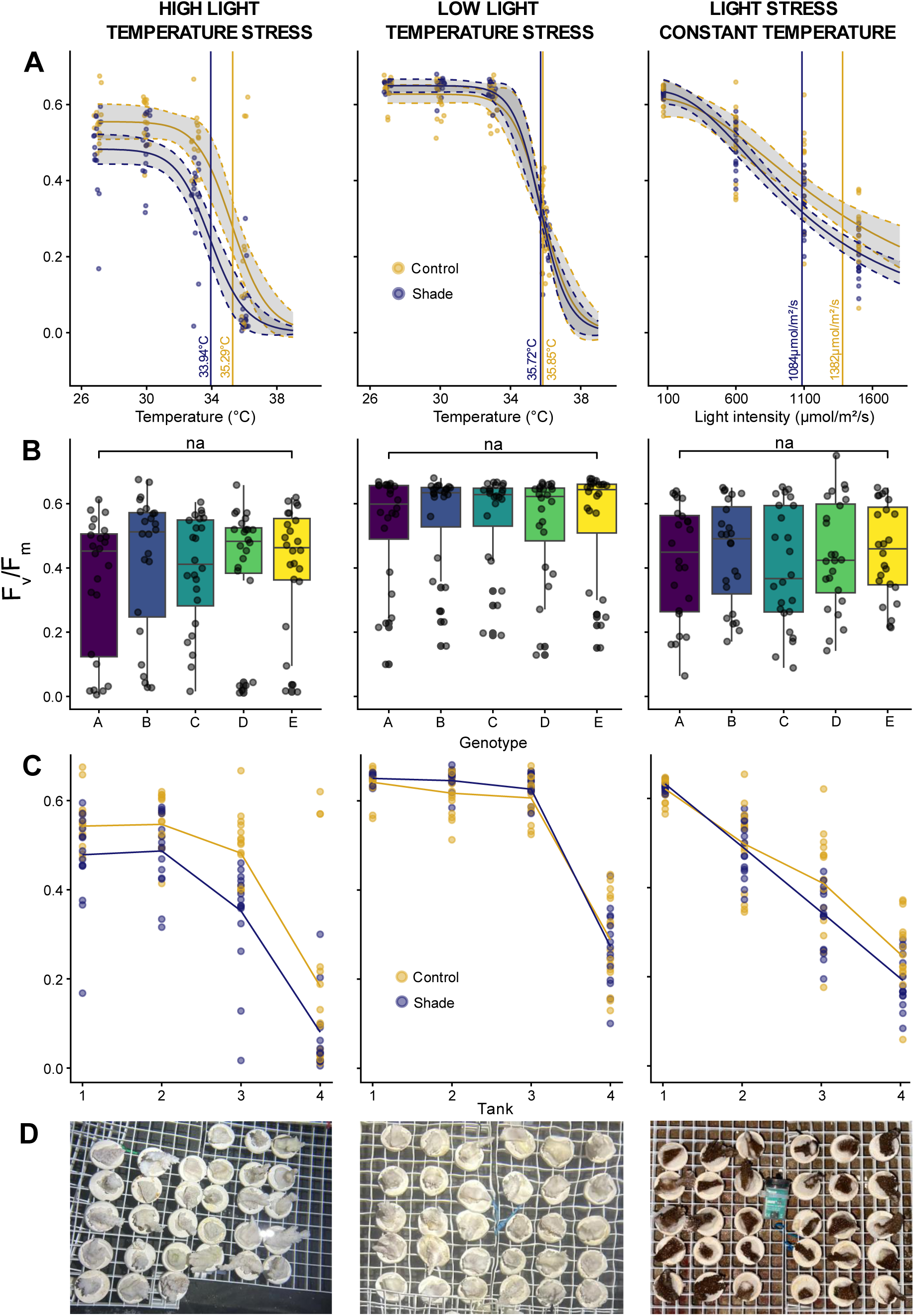
A) Corals photosynthetic efficiency for each CBASS assay and each light condition. Log-logistic dose-response curves were fitted to the PAM (F_v_/F_m_) data, with dashed lines depicting their 95% confidence intervals. Vertical lines represent the ED50 for each historical light condition. B) Boxplot of genotypic variation across each CBASS assay. C) Random intercept mixed models depicting the influence of temperature and light intensity on the observed data. Dots represent raw data, while solid lines show model predictions for each CBASS assay. D) Post-CBASS visual bleaching assessment of corals exposed in the most severe tank of each assay (Tank 4)

### Fluorometry

The dark-acclimated photosynthetic efficiency of each coral nubbin was measured using a MINI-PAM II (Pulse-Amplitude Modulated, WALZ, Germany) with settings matching the manufacturer’s instructions (MI: 3, Gain: 6, DAMP: 2, SI: 8, SW: 0.8 s). The measured F_v_/F_m_ ratio of PSII was used to compare the dark-adapted, pre-photosynthetic fluorescent state (F_0_) to maximum fluorescence (F_m_) for each coral. F_v_/F_m_ is commonly used as an indicator of algal performance under temperature and light stress (Lichtenthaler et al. 2005). Low F_v_/F_m_ values would indicate a state close to photo saturation, while higher values would be found under optimal conditions. We measured this non-destructive stress proxy in the dark at 8pm after each CBASS assay (Evensen et al. 2021). In addition to measuring maximum quantum yield, we photographed coral nubbins of each tank at the end of each assay to obtain a visual coral bleaching assessment. Coral bleaching was analyzed as a three-level factor variable, with values being “healthy”, “pale”, and “bleached”.

### Statistical analyses

All statistical analyses and data plotting were conducted in Rstudio v.2024.04.2+764 (R Core Team), with the ‘ggplot2’ package used to plot figures and the Inkscape program used to polish figures (Bah 2011). Further, we use the ‘drc’ R package to fit log-logistic dose-response curves to the F_v_/F_m_ data. These curves were used to determine the effective dose 50 (ED50) for both control and shade corals across each assay. ED50s correspond to the temperature or light values for which the F_v_/F_m_ response is lowered by 50%, and is therefore used as an empirically derived proxy for stress tolerance (Evensen et al. 2022). ED50s were hereafter compared among light treatments through a one-way Analysis of Variance (one-way ANOVA) to test for the effect of coral light history on stress response. Lastly, we built generalized linear mixed models using the textitlme4() function of Rstudio to assess the relative influence of temperature and light intensity on the F_v_/F_m_ data, with temperature and light variables scaled using the *scale()* function, *Genotype* as random effect, and *Treatment* as a random variable nested within *Assay*. Aikake’s Information Criterion (AIC) was used as an indicator of model performance.

## Results

### Nursery experiment

Corals were successfully grown for two years at the HIMB coral nursery via the above-described setup. Corals in the control treatment received an average photosynthetic solar irradiance of 176.8µmol.m^-^².sec^-1^, with daily maxima ranging from 370 to 1939µmol.m^-^².sec^-1^. Average solar irradiance in the shade treatment was 25.0µmol.m^-^².sec^-1^, with daily maxima ranging from 35.2 to 240.5µmol.m^-^².sec^-1^ (Supplementary Fig. S1). Irradiance data converted to watts.m^-2^ as recommended by Tagliafico et al. (2022) are available in the Supplementary Materials. Sea surface temperature ranged between 21.7°C and 27.9°C, with a mean value of 24.8°C. Five genotypes did not show any sign of mortality, while three others experienced moderate to severe mortality in some fragments. However, the number of fragments that were completely dead did not differ significantly among light conditions (n = 5 in the control treatment and n = 3 in the shade treatment). No extreme weather event occurred within these two years, and none of these corals experienced natural bleaching during this two-year period.

### Physiology

Prior to the application of shade, we detected no differences in ΔF/F_m_’ across light treatments (ANOVA, *p*-value = 0.86). However, after the application of shade, ΔF/F_m_’ values started to diverge between treatments (Fig. 2A). From the third sampling day onwards, average ΔF/F_m_’ became significantly higher in the shade treatment than in the control (ANOVAs, *p*-values < 0.01), with values ranging from 0.455 to 0.712 under shaded light, and from 0.154 to 0.556 at ambient light. The observed daily variations were likely due to changes in cloud cover and photosynthetically active radiation. Similarly, symbiont density was significantly higher in the shade than in the control (Fig. 2B; ANOVA, *p*-value < 0.01), with corals from the control treatment having nearly two times lower symbiont density than the colonies growing in the shade. The log-log scale was used to establish a linear relationship between growth rate and initial buoyant weight. We found growth rate to be significantly higher in the control treatment compared to the shade regardless of initial fragment size (Fig. 2C; ANOVA, *p*-value < 0.01), ranging from increases in buoyant weight of 8.7% to 91.2% in the control and from 6.5% to 55.9% in the shade.

### Effective Doses 50 (ED50s)

For each of the three CBASS assays, a dose-response curve was fitted to the PAM (F_v_/F_m_) data, from which ED50s were calculated. After temperature stress under high light, the ED50 of the control corals was higher than the ED50 of the shaded corals (35.29°C and 33.94°C respectively, Fig. 3A). This nearly 1.5°C difference in ED50 was found to be significant (one-way ANOVA, *p*-value < 0.01) and suggests control corals performed better than shaded colonies during this assay. However, we found no significant difference in ED50 when the same heat treatment was applied at low light (Fig. 3B, one-way ANOVA, *p*-value = 0.32), indicating that long-term shading did not seem to lower heat tolerance. Additionally, we found a reduction of the decline of photosynthetic efficiency in the low-light assay regardless of historical light conditions. Results indicate a significant increase of ED50s for both control (one-way ANOVA, *p*-value < 0.04) and shaded colonies (one-way ANOVA, *p*-value < 0.01) between the high-light assay and the low-light assay. ED50s increased from 35.29°C to 35.85°C for corals that grew in ambient light, and from 33.94°C to 35.72°C for corals that grew under shade. Lastly, we ran a CBASS with variable light stress at constant water temperature (Fig. 3B) and found corals grown in ambient light conditions had higher photosynthetic efficiency than those grown under shade (one-way ANOVA, *p*-value <0.01). Here, the distribution of F_v_/F_m_ was also relatively evenly spread, with control corals having an ED50 of 1382 µmol.m^-2^.sec^-1^, against 1084 µmol.m^-2^.sec^-1^ for shaded corals.

### Post-CBASS bleaching

Overall, we found no variability in bleaching response within the fragments of each tank, regardless of the temperature by light treatment. After the heat stress assay at ambient light, all coral fragments from Tank 4 were completely bleached, corals from Tank 3 were pale (Fig. 3D), and corals from Tank 1 and Tank 2 did not show any sign of bleaching. Identical observations were made after the heat-stress assay at low light. However, in the light-stress treatment, no bleaching or paling was observed in any fragment regardless of tank, despite an equivalently sharp drop in F_v_/F_m_ as in the temperature stress assays (Fig. 4).

**Fig. 4:**
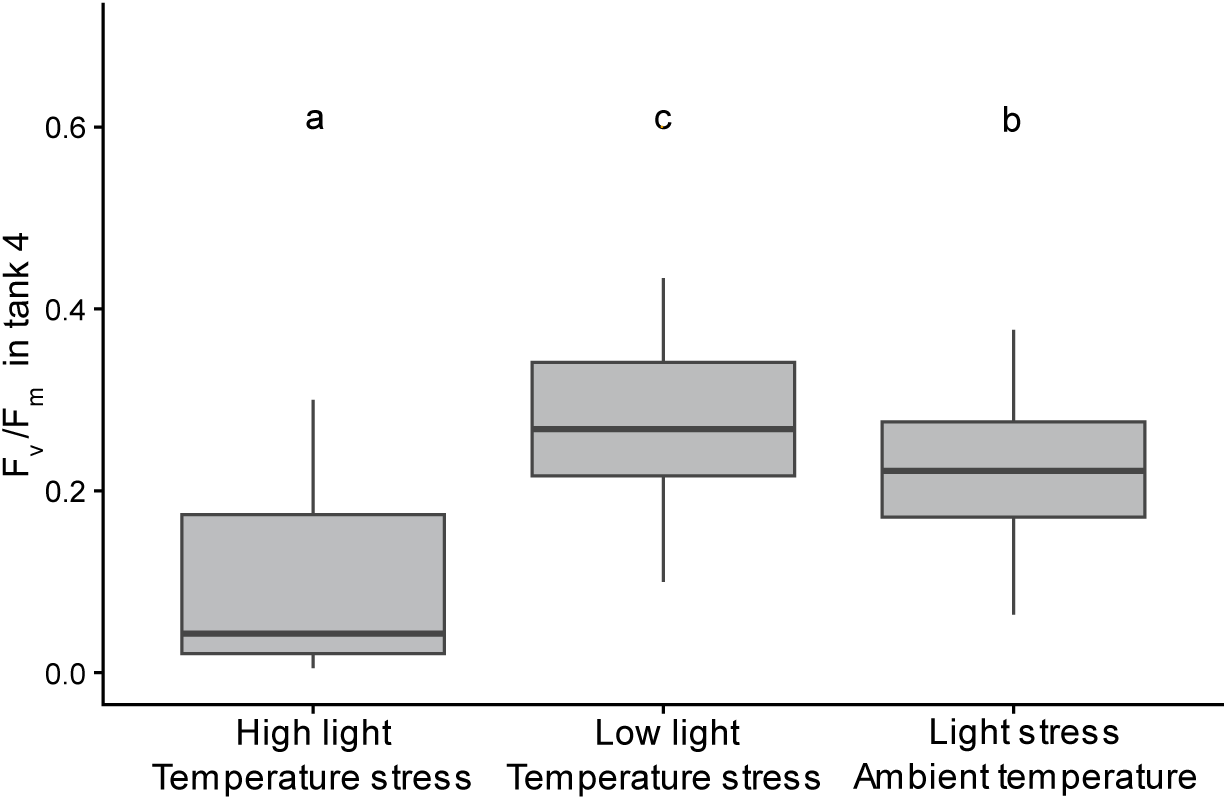
Boxplots of F_v_/F_m_ values of Tank 4 for each CBASS assay. Letters indicate significance levels

### Variation in F_v_/F_m_

We found that average F_v_/F_m_ values in Tank 4 (i.e. the tank corresponding to the most severe stress) differed significantly between each assay, with the high-light, heat-stressed nubbins having the lowest F_v_/F_m_, followed by the light-stressed nubbins, and by the low-light, heat-stressed nubbins (Fig. 4, *p*-value < 0.01). To dig deeper into the underlying processes that drive F_v_/F_m_ variation, we plotted the obtained F_v_/F_m_ values against dose values (i.e. temperature or light values) for each assay (Fig. 3C) using random intercept mixed models with *Treatment* as a random variable nested within *Assay*. Essentially, light and temperature were both found as significant predictors of F_v_/F_m_ (Mixed model, *p*-value < 0.01), with light intensity having roughly twice the explanatory power of temperature (F-value_light_ = 25.59; F-value_temperature_ = 13.2). Interestingly, the interaction of light and temperature was not a significant predictor of F_v_/F_m_ (*p*-value = 0.83). We did not include the *Genotype* variable in the final model, as we found it to be a very poor, non-significant predictor of F_v_/F_m_ (p.value = 0.52). This was also corroborated by the fact that we did not detect any genotypic difference in F_v_/F_m_ response within any assay (Fig. 3B).

## Discussion

### Light acclimation

In this study, we show that the light history of *Montipora capitata* coral fragments originating from the same genotype can lead to drastically different responses to light stress. Acclimation to decreased light intensity entailed higher photosynthetic efficiencies and symbiont densities, as well as slower growth rates (Fig. 2). The ΔF/F_m_’ time-series suggests that this acclimation occurred over the course of the first three days, as ΔF/F_m_’ values between control and shade started to diverge from one another as soon as we applied shade and remained constant from the third sampling day onwards. The slight drop in ΔF/F_m_’ after two days in the control treatment was explained by an increase in solar radiation (Supplementary Fig. S1). This drop was however not present in the shaded corals, where the photosynthetic rate increased instead. This suggests that shading may have protected these colonies from the light stress the control corals experienced.

Symbiont densities in the control treatment were lower than in the shaded one following six months of exposure, as daily metabolic requirements are met more easily when light is not limited (Fig. 2B; Mason et al. 2020). This also explains why we found significantly higher growth rates under ambient light relative to the shade treatment regardless of the initial size of our experimental fragments (Fig. 2C). As light was not the limiting factor in the control treatment, corals could grow more rapidly, whereas shaded corals appeared down-regulated. This suggests a trade-off between protection to high levels of solar irradiance and growth rate, i.e., a balance between the coral experiencing photodamage and being light-limited (López-Lodoño et al. 2022). Fast-growing corals exposed to high levels of irradiance may be experiencing photodamage, whereas corals at decreased light intensity are less exposed to photodamage but grow slower. These results also highlight the phenotypic plasticity of *M. capitata* at the physiological level in response to different levels of light intensity, as already described for micro-morphological (Bhagooli 2003) and reproductive traits (Henley et al. 2021) as well as for gene expression (Drury et al. 2022). Further research at the molecular or cellular level will be needed to gain insight into the processes that drive this plasticity.

### Stress responses

During the heat-stress assay at high light, we found a significant difference in ED50 between control and shaded colonies (Fig. 3A). This difference was not observed under low light, so we conclude that historical light conditions had no effect on photosynthetic efficiency under high temperatures (Fig. 3A). This demonstrates that long-term shading does not lower coral heat tolerance. Instead, the observed difference of ED50s in the first assay is likely due to light stress. Corals with a history of growing under shade were acclimatized to levels of light intensity of a much lower range than the 600µmol.m^-^².sec^-1^ used during the high-light temperature stress assay.

Another interesting pattern between the high-light and low-light temperature stress assays is the greater variability in F_v_/F_m_ in high light. Even at 27°C, the lowest temperature treatment, some genotypes, particularly from the shade treatment, showed a reduced photosynthetic efficiency at high light levels. On the other hand, F_v_/F_m_ values were higher and more homogeneous in the heat stress assay at low light (Fig. 3A, 3C). This suggests that photo-stress may already occur at 600µmol.m^-^².sec^-1^ for corals that are low-light acclimated. This reflects the importance of considering light acclimation prior to using CBASS, as underestimated differences in the light history of different coral fragments may lead to a biased interpretation of the ED50s. This supports evidence that the irradiance exposure of corals during any thermal stress experiments is critical and can have significant impacts on measures of symbiont health (Lesser 2024). The light-stress assay performed at ambient temperature also confirms that corals exposed to ambient light conditions are more resistant to light stress than those grown under shade (Fig. 3).

ED50s increased significantly when light intensity was lowered during heat stress (Fig. 3A). This supports the hypothesis that a reduction of light intensity can mitigate stress (Baird et al. 2016; Coelho et al. 2017; Butcherine et al. 2023; Winslow et al. 2024), and is consistent with the idea that light and temperature have synergistic effects on coral health (Fitt et al. 2001). However, we found no differences in post-assay bleaching susceptibility between the heat-stress assays at ambient light and at low light (Fig. 3D). This is probably explained by the fact that CBASS is designed to go far beyond tolerance thresholds, and is meant to force bleaching to happen even in extremely tolerant corals. Corals, outside of extreme environments, generally do not experience such rapid, extreme increases in ocean temperatures, and the extent to which post-CBASS bleaching observations can be extrapolated to natural conditions is yet to be explored. Further research aiming at comparing CBASS with standardized, long-term bleaching experiments (Grottoli et al. 2021; Evensen et al. 2023; Klepac et al. 2024) or in-situ observations would allow better understanding of the extent to which shading mitigates coral bleaching.

### Different stress pathways

Elevated temperature and light are both environmental stressors that can drastically affect coral health, eventually leading to coral bleaching (Coles & Jokiel, 1978). The severity of the stress is negatively correlated with the maximum quantum yield of PSII, commonly referred to as F_v_/F_m_. Therefore, this metric is commonly used as a proxy for bleaching severity (Evensen et al. 2021; Klepac et al. 2024). Surprisingly, the most linear relationship between F_v_/F_m_ and stress severity was obtained during our light stress assay, where temperature was held at MMM, and yet coral bleaching was not observed (Fig. 3C & 3D). Alternatively, heat stress severity showed a non-linear relationship with F_v_/F_m_ in the first two heat stress assays, where bleaching was observed in Tank 4 (Fig. 3C & 3D). It is important to note that we found significantly lower F_v_/F_m_ values in Tank 4 of the light-stress assay, where corals did not bleach, compared to the low-light heat stress assay, where corals in Tank 4 did bleach (Fig. 3D; Fig. 4). Across all assays, each temperature or light dose in Tank 4 was therefore strong enough to induce an ecologically significant, sharp drop in F_v_/F_m_, but this did not necessarily lead to bleaching. As light stress triggers damage in the symbiont’s photosystems (Oakley & Davy 2018; Helgoe et al. 2024), this work may suggest that F_v_/F_m_ is a predictor of oxidative stress in the endosymbionts rather than of cell destruction or symbiont expulsion pathways. The corals in the 36°C treatment tanks may have experienced the Unfolded Protein Response cascade due to the exceptionally high and acute temperatures, which led to coral cell necrosis/apoptosis in these fragments (Helgoe et al. 2024). This is consistent with findings from Voolstra et al. (2020), who recorded sharp declines in host protein concentrations in corals in CBASS assays but only in the Tank 4 treatment. Alternatively, corals in the 1600µmol.m^-^².sec^-1^ tank, which also recorded equivalently low F_v_/F_m_, may have experienced photoinhibition, recovery from which occurred during night-time protein repair, as the proteins themselves were not damaged by heat stress (Helgoe et al. 2024). Therefore, these corals did not visually bleach or die. This highlights the importance of elucidating the different pathways that lead to bleaching, and questions the relevance of measuring the maximum quantum yield of PSII when assessing thermal bleaching response using CBASS. This study supports the work of Klepac et al. (2024), who also found that F_v_/F_m_ response and bleaching response were not necessarily linked during their CBASS experiment. We therefore encourage combining PAM data with transcriptomics (Savary et al. 2021; Drury et al. 2022) or host protein-content analyses (Voolstra et al. 2020) to accurately determine the causes of bleaching in future heat or light stress experiments.

### Implications for conservation

Coral reefs are solar-driven ecosystems whose species and functional diversities strongly vary with depth, and therefore light availability (Roberts et al. 2019; Tamir et al. 2019; López-Lodoño et al. 2022). Differences in solar irradiance may result in shifts in species composition, as well as in photophysiology or light tolerance. From a conservation standpoint, we recommend to gradually light-acclimate coral colonies when moved from a low-light area to higher-light areas, especially if rearing of coral larvae was done at different light regimes (Hancock et al. 2021; Ransby et al. 2023) or in land-based nursery with darker ambient light intensity (Gantt et al. 2023). Optimal solar irradiance at outplant sites should also be investigated, as light niches are species-specific (Roberts et al. 2019; Winslow et al. 2024) and affected by site-specific water optical properties (López-Lodoño et al. 2022). Further, if using the CBASS to identify heat-tolerant individuals, it is essential to ensure they are collected from the same light environment or are sufficiently light-acclimated to avoid erroneous conclusions biased by photoacclimation. Additionally, in our experiment long-term shading did not lower corals heat tolerance, while short-term shading reduced the decline of photosynthetic efficiency. This suggests that both short-term and long-term shading strategies have no negative impact on the survival of the shaded corals while protecting them from bleaching, and are therefore helpful approaches to mitigate coral bleaching in the field.

