## Supplementary Information for "Shading does not lower thermal tolerance in the coral *Montipora capitata*: implications for conservation intervention"

**Supplementary material for the research article entitled “Shading does not lower thermal tolerance in the coral *Montipora capitata*: implications for conservation intervention”.**

### Nursery experiment

We have converted the irradiance data from the long-term experiment from µmol.m².sec$^{-1}$ into watts m$^{-2}$ here for future reference for the research community, using a x0.217391304 conversion factor as recommended by Tagliafico et al. (2022). Corals in the control treatment received an average photosynthetic solar irradiance of 38.44 watts m$^{-2}$, with daily maxima ranging from 80.435 to 421.52 watts m$^{-2}$. Average solar irradiance in the shade treatment was 5.439 watts m$^{-2}$, with daily maxima ranging from 7.654 to 52.283 watts m$^{-2}$.

### Supplementary figures

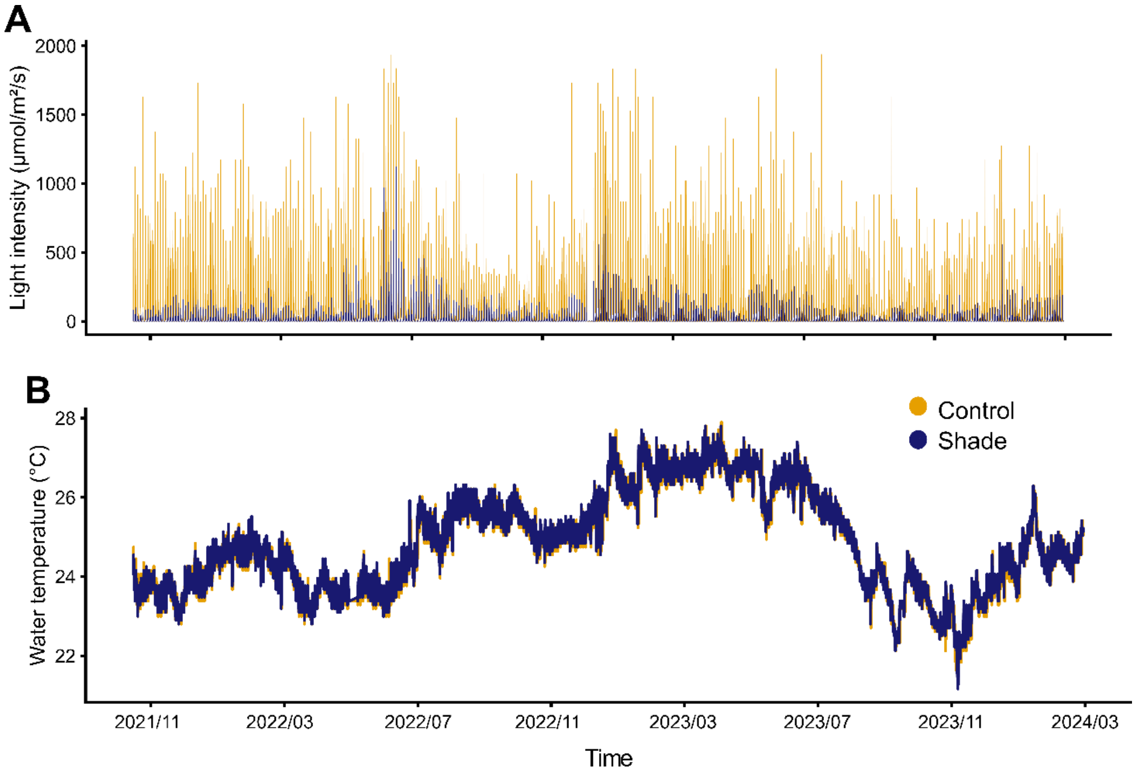

Supplementary Fig. S1: A) Recorded light intensity (µmol.m².sec$^{-1}$) and B) water temperature (°C) at the HIMB coral nursery during the two years of growth.

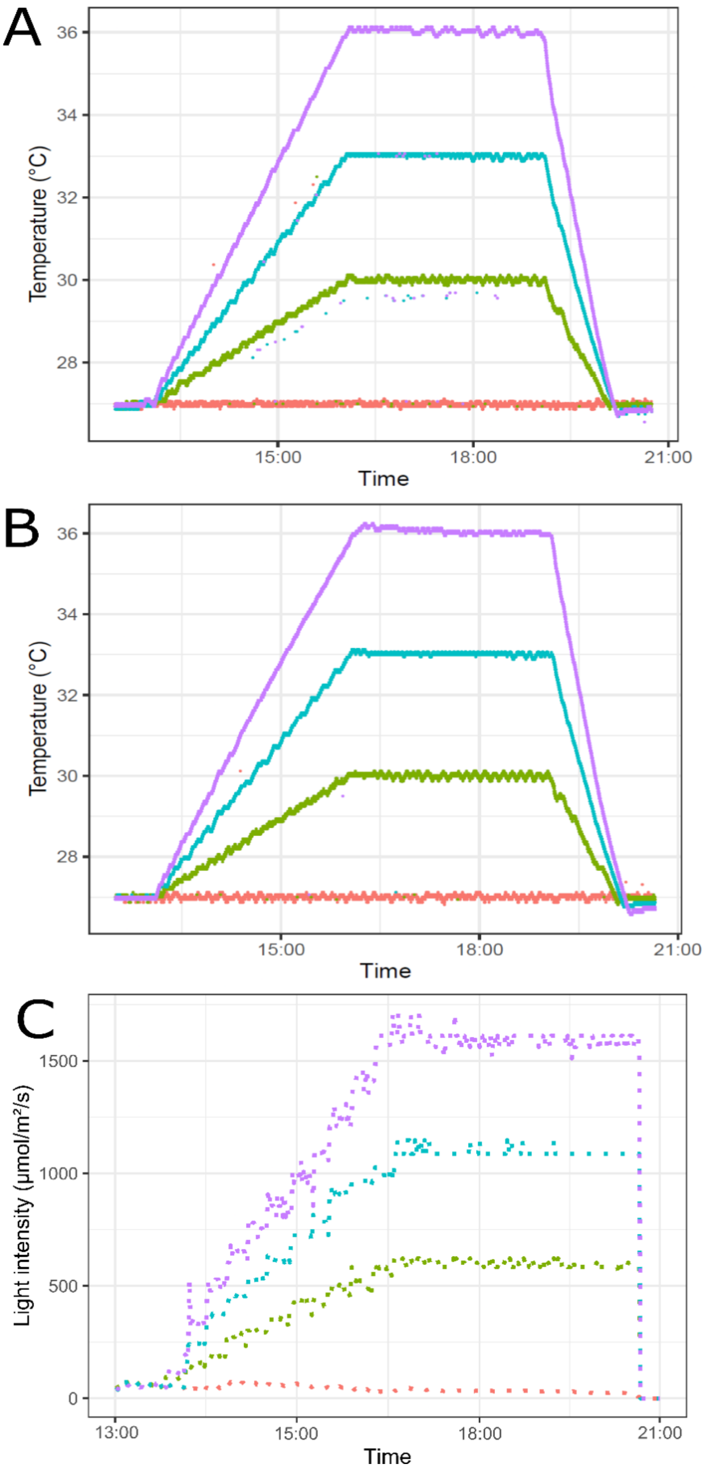


Supplementary Fig. S2: Temperature and light profiles of each CBASS assay. A) High-light temperature stress assay. B) Low-light temperature stress assay. Data for these two assays were retrieved via the log files of heaters and chillers controlled via the Arduino platform. C) Light stress assay at constant temperature. Data for this assay was retrieved from HOBO pendant loggers placed in each tank during the assay.
